# SiPLATZ12 transcript factor regulates multiple yield traits and salt tolerance in foxtail millet (*Setaria italica*)

**DOI:** 10.1101/2022.07.01.498439

**Authors:** Shenghui Xiao, Yiman Wan, Linlin Zhang, Sha Tang, Yi Sui, Yichao Bai, Yan Wang, Miao Liu, Jiayin Fan, Shizhong Zhang, Jinguang Huang, Guodong Yang, Kang Yan, Xianmin Diao, Chengchao Zheng, Changai Wu

## Abstract

Grain yield and salt tolerance are critical for crop production. However, the genetic and biochemical basis underlying the trade-off of these characters remain poorly described in crops. We show here that SiPLATZ12 transcription factor positively regulates multiple elite yield traits at the expense of salt tolerance in foxtail millet. SiPLATZ12 overexpression increases seed size, panicle length, and stem diameter, while reduces plant height and salt tolerance of foxtail millet. A 9-bp insertion in the *SiPLATZ12* promoter has significant effects on the different expression of *SiPLATZ12*, multiple yield traits, and salt tolerance between foxtail millet and its wild ancestor, green foxtail. Moreover, SiPLATZ12 upregulates the expression of genes involved in seed development, but repressing the transcription of most *NHX, SOS*, and *CBL* genes to regulate Na^+^, K^+^ and pH homeostasis. Therefore, our results uncover a domesticated site that could be used to improve grain yield and salt tolerance in foxtail millet.

## Introduction

Foxtail millet (*Setaria italica*), one of the most ancient domesticated crops in the world, is mainly cultivated in arid and semi-arid regions in Asia, Eurasia, and Africa for food, feed, fuel, and bioenergy (Brutnell *et al*., 2010; Jia *et al*., 2013; Yang *et al*., 2020; Zhang *et al*., 2012). There is a demand for new foxtail millet cultivars with high yield and strong abiotic stress tolerance. However, the yield and salt tolerance of foxtail millet are inherited quantitatively, so increasing yield and salt tolerance through traditional genetic improvement is difficult and time consuming. Once the key genes controlling important agronomic traits and their interrelationship are identified, improving foxtail millet through biotechnology will be very effective.

To achieve high yield, plants require not only high yield component traits, but also an optimal architecture that, to a large extent, is determined by plant height. Moreover, heading date (or flowering time) is a critical determinant enabling crops to adapt to seasonal changes and make maximum use of the temperature and sunlight. Yield per plant in foxtail millet is determined by two component traits: number of grains per panicle (panicle size) and grain weight (grain size). Multiple genetic factors controlled seed size have been revealed in Arabidopsis. These include the VQ domain containing protein HAIKU1 (IKU1), the leucine rich repeat receptor kinase IKU2, MINI-seed3 (MINI3), short Hypocotyl under Blue 1 (SHB1), the KLUH (KLU), fertilization independent seed 2 (FLS2), and cytokinin independent 1 (CKI1) coding genes (Kesavan *et al*., 2013; Li *et al*., 2016; Luo *et al*., 2005; Wang *et al*., 2010; Zhou *et al*., 2009). In rice, serine carboxypeptidase (GS5) (Li *et al*., 2011), gibberellic acid-stimulated regulator (GASR)7/grain weight (GW6) (Shi *et al*., 2020; Tang *et al*., 2021), and OsALMT7 (Heng *et al*., 2018) have been identified to function in seed size control. Besides, dwarf/semidwarf (D/SD) and stem diameter also contribute to high grain yield mainly by increasing the lodging-resistance of crops. Molecular cloning and functional analyses of several genes associated with plant height in rice and wheat have shown that these genes are mostly related to the synthesis and regulation of the phytohormone gibberellin (Peng *et al*., 1999; Spielmeyer *et al*., 2002). Widespread utilization of such mutant gene, *sd1*, in worldwide breeding programs of staple crops, such as rice and wheat, results in the development of modern semidwarf lodging resistant cultivars (Monna *et al*., 2002; Spielmeyer *et al*., 2002). Extensive studies in *Arabidopsis thaliana* have established that flowering time is controlled by multiple pathways (Siplson *et al*., 2002), of which the photoperiod pathway acts as a major one. A key component functioning in this pathway is the CONSTANS (CO) TF. One of the downstream targets of CO is flowering locus T (FT), which controls flowering time by integrating input from various pathway (Santiag *et al*., 2021). Although the function of most of these genes has not been characterized, Loose Panicle 1, a group I WRKY transcription factor (TF), has been revealed to control plant height, panicle and seed size in foxtail millet (Xiang *et al*., 2017). Besides, SiMADS34 has been recently characterized to regulate panicle architecture, grain weight, and seed shape in foxtail millet (Hussin *et al*., 2021).

Except for plant architecture, external abiotic stresses seriously threat the food security worldwide. Salt stress is one of the main abiotic stresses. The salt tolerance of plants has been proved to be largely associated with cytoplasm Na^+^ homeostasis which is maintained by membrane localized Na^+^/H^+^ antiporters (NHXs) and their regulatory proteins, such as Salt Overly Sensitives (SOSs) and Calcineurin B-Like proteins (CBLs) (Halfter *et al*., 2000; Liu and Zhu, 1998; Quintero *et al*., 2011; Zhou *et al*., 2022). The transporter activities of NHX7/SOS1 are activated by SOS2-SOS3 (CBL4)/ SCaBP8(CBL10) complex (Halfter *et al*., 2000; Qiu *et al*., 2002; Quan *et al*., 2007). Studies also showed that the NHX proteins are important for compartmentalization of K^+^ into vacuoles, cellular pH homeostasis (Barragán *et al*., 2012; Bassil *et al*., 2011a; Bassil *et al*., 2011b), and protein trafficking (Reguera *et al*., 2015). Thus, NHX operations play multiple roles in stomatal regulation (Barragán *et al*., 2012), plant growth (Bassil *et al*., 2012; Bassil *et al*., 2011a; Bassil *et al*., 2011b), and silique and seed development (Wu *et al*., 2016). However, the function of most of these genes has been poorly investigated in foxtail millet.

PLATZ TFs are a class of plant-specific zinc-dependent and A/T-rich sequence binding proteins (Nagano *et al*., 2001). In *Arabidopsis*, PLATZ1 positively regulates drought tolerance in vegetative tissues (Gonzalez-Morales *et al*., 2016). AtPLATZ2 negatively regulates plant salt tolerance (Liu *et al*., 2020). ORESARA15, as known as AtPLATZ3, promotes leaf cell proliferation during earlier stages of development and suppresses leaf senescence during later stages (Kim *et al*., 2018). In crops, PLATZs from maize (*Zea mays*) (Li *et al*., 2017), rice (*Oryza sativa*) and wheat (Wang *et al*., 2019; Zhou and Xue, 2020; Guo *et al*., 2022) regulate endosperm development and seed filling, grain length and number, respectively. PLATZs therefore play important roles in plant growth, development and abiotic stress responses. However, the function of PLATZs has not been characterized in foxtail millet.

Here, we identified SiPLATZ12 transcription factor as a positive regulator of multiple yield traits but a negative regulator of salt tolerance in foxtail millet. A 9-bp insertion in the *SiPLATZ12* promoter significantly associated with the function of *SiPLATZ12*. Moreover, SiPLATZ12 regulates the expression of *IKU1, IKU2, MINI3, SHB*1, *KLU, FLS2*, and *CKI1*genes for seed development and *SiNHX, SiSOS*, and *SiCBL*s genes for both development and salt tolerance. In particular, SiPLATZ12 directly binds to the A/T rich sequences in *SiNHX2, SiCBL4* and *SiCBL7* promoters. This study provides genetic resources for breeding high yield cultivars of foxtail millet by increasing *SiPLATZ12* expression or salt tolerant cultivars by editing *SiPLATZ12*.

## Results

### SiPLATZ12 affects multiple yield traits and salt tolerance in foxtail millet

To identify *PLATZ* genes in foxtail millet, we scanned the ‘Yugu1’ genome assembly for open reading frames (ORFs) containing the PLATZ domain sequences from *Arabidopsis thaliana*. This identified 16 *PLATZ* genes in foxtail millet, which were named *SiPLATZ1* through *SiPLATZ16* on the basis of chromosomal localization (Table S1). Neighbor-joining analysis classified SiPLATZs and PLATZs from maize, rice, and *Arabidopsis* into three subfamilies (Appendix Fig S1A). The subfamily homologous to maize FL3, including *SiPLATZ*7-*13*, was selected for further analysis.

To quantify the expression patterns of *SiPLATZ*7-*13*, we carried out quantitative reverse transcription-PCR (RT-qPCR) and found that except for *SiPLATZ10*, other *SiPLATZ* genes of this subfamily exhibited higher transcript levels in roots, followed by in panicles and shoots, than that in leaves (Appendix Fig S1B). *SiPLATZ8* and *SiPLATZ12* showed the highest transcript levels in roots of 14-day-old seedlings. Moreover, these *SiPLATZ* genes were also induced at 3 and 24 h after salt, alkaline, temperature and ABA treatments (Appendix Fig S1C).

To characterize the function of *SiPLATZ12*, we generated transgenic foxtail millet lines in the ‘Ci846’ background. Transcript levels of *SiPLATZ12* in 14 independent 35S::SiPLATZ12 transgenic lines were verified by RT-qPCR (Appendix Fig S2A), and three lines (#22, #24, #28) were selected for further investigation. The transgenic plants exhibited significantly larger seeds with up to 29.7% longer and 36.5% wider seeds, resulting in 40.3% higher TGW than those of ‘Ci846’ (Fig 1A-D). The transgenic plants also exhibited 32.2% longer and 36.7% wider panicles (Fig 1E-G), 17.7% lower grain number per panicle, and 32.2% higher grain weight per panicle (Fig 1H and I) than those of ‘Ci846’. Moreover, compared with ‘Ci846’, the three transgenic plants also showed increased main stem diameter (Fig 1J and K), reduced plant height and leaf number (Fig 1L-N), as well as delayed heading time (Fig 1L). The higher transcript levels of *SiPLATZ12* in panicles, especially young panicles than other indicated tissues, such as flag leaves, roots, stems and leaves of young seedlings confirmed the role of *SiPLATZ12* in regulating the corresponding organ development (Appendix Fig S2B). These results indicated the function of *SiPLATZ12* in positively regulating multiple elite yield traits in foxtail millet.

**Figure 1.**
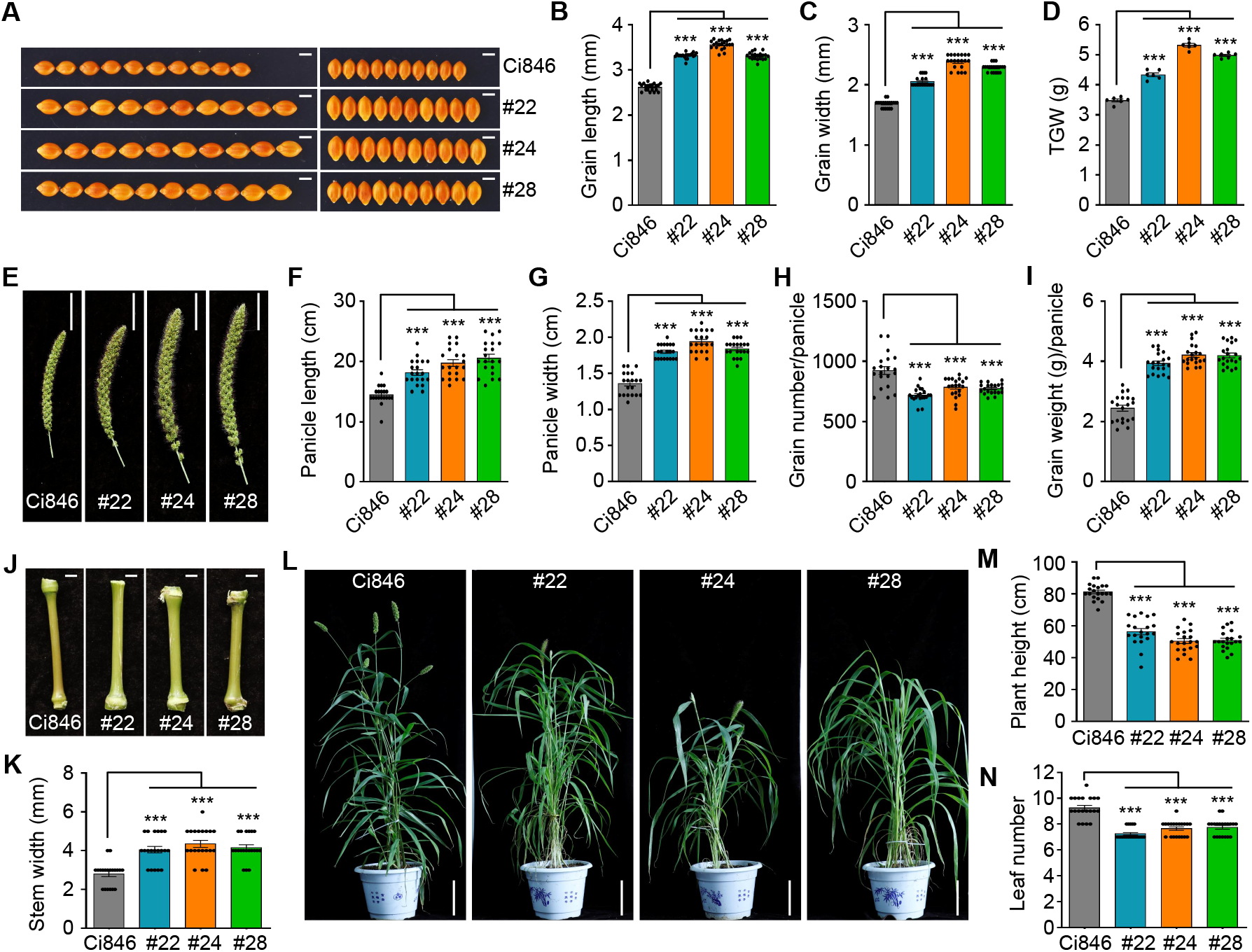
SiPLATZ12 affects multiple yield traits in foxtail millet. A Seed shape of *35S::SiPLATZ12* transgenic plants and ‘Ci846’ (the background material) under normal conditions. Scale bars = 2 mm. B-D Seed length (B), width (C) and TGW (D) of ‘Ci846’ and *35S::SiPLATZ12* transgenic plants. At least 30 seeds were measured in each replicate. Three independent replicates were conducted. Values are given as the mean ± SD, \*\*\**P* < 0.001 by Student′s *t*-test. E Panicle shape of ‘Ci846’ and *35S::SiPLATZ12* transgenic plants under normal conditions. Scale bars = 5 cm. F-I Panicle length (F) and width (G), grain number (H) and weight (I) per panicle of ‘Ci846’ and *35S::SiPLATZ12* transgenic plants. At least 20 panicles were measured in each replicate. Three independent replicates were conducted. Data are represented as the mean ± SD, \*\*\**P* < 0.001 by Student′s *t*-test. J, K Stem phenotype (J) and diameter (K) of ‘Ci846’ and *35S::SiPLATZ12* transgenic plants. Scale bars = 4 mm. At least 20 stems were measured in each replicate. Three independent replicates were conducted. \*\*\**P* < 0.001 by Student′s *t*-test. L *35S::SiPLATZ12* transgenic foxtail millet grew slower and heading later than ‘Ci846’ under normal conditions. Scale bars = 10 cm. M, N Plant height (M) and leaf number (N) of ‘Ci846’ and *35S::SiPLATZ12* transgenic plants. At least 20 plants were used in each replicate. Three independent replicates were conducted. \*\**P* < 0.01 and \*\*\**P* < 0.001 by Student′s *t*-test.

Moreover, when 3-day-old seedlings were transferred into Hoagland medium containing series of concentrations of NaCl, growth of transgenic seedlings was seriously inhibited, with 40%–50% shorter roots and 30% lower fresh weight than ‘Ci846’ under 250 mM NaCl (Fig 2). These results illustrated that *S*i*PLATZ12* negatively regulate salt tolerance in foxtail millet. However, the longer roots and larger shoots of *SiPLATZ12* transgenic foxtail millet than ‘Ci846’ under normal conditions and low concentrations NaCl (< 100 mM) (Fig 2A) might be due to their large seeds.

**Figure 2.**
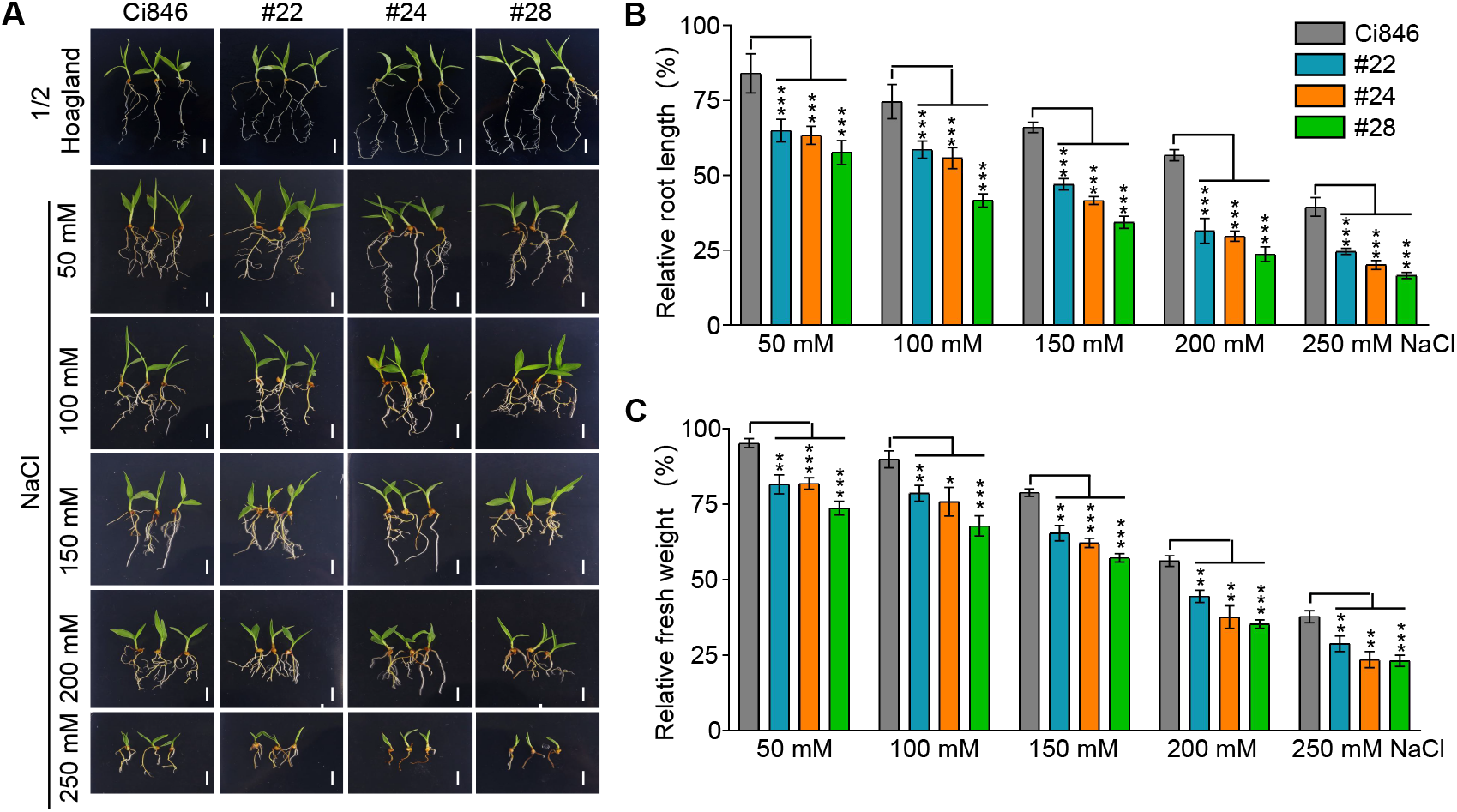
SiPLATZ12 negatively regulates salt tolerance in foxtail millet. A Phenotypes of seven-day old Ci846’ and *35S::SiPLATZ12* transgenic seedlings under normal and salt stress conditions. Scale bars = 1 cm. Seedlings were grown in half-strength Hoagland solution containing a series of indicated concentrations of NaCl. B, C Relative root length (B) and fresh weight (C) of seven-day old Ci846’ and *35S::SiPLATZ12* transgenic seedlings under normal and salt stress conditions. At least 20 seedlings were used in each replicate. Three independent replicates were conducted. \**P* < 0.05, \*\**P* < 0.01 and \*\*\**P* < 0.001 by Student′s *t*-test.

In addition, the function of *SiPLATZ12* was verified by heterologously expressing in Col-0 *Arabidopsis*. The transgenic Arabidopsis lines (#1, #4 and #6) were confirmed by RT-qPCR (Appendix Fig S3A). Consistent with the results in foxtail miilet, the *SiPLATZ12* transgenic *Arabidopsis* also showed larger seeds and higher TGW than those of the WT (Appendix Fig S3B and C). Besides, these transgenic *Arabidopsis* seedlings also displayed slower growth and lower fresh weight than the WT (Appendix Fig S3E and F). The salt tolerance of these transgenic *Arabidopsis* seedlings were reduced obviously with lower germination rate, growth, fresh weight and SUR compared with the WT (Appendix Fig S3D-G). These results confirmed the positive role of *SiPLATZ12* in regulating TGW but negative role in regulating salt tolerance in plants.

### A 9-bp insertion in the *SiPLATZ12* promoter improves its expression and yield traits in millets

To verify the function of *SiPLATZ12* genetically, we compared the nucleotide differences of DNA regions including 3 Kb upstream and 2 Kb downstream and gene body of *SiPLATZ12* among 1893 millet accessions. Total of 18 variations, representing 158 haplotypes, exist within this region (Fig 3A). Five major haplotypes with high frequency (more than 40 millet accessions each) were selected for further analysis (Fig 3B and C). Hap1 was the predominant haplotype, with 1099 foxtail millet (cultivated) accessions. Hap18 was the second major haplotype, with 264 green foxtail accessions (wild varieties). Hap2 and Hap14 were represented by 77 and 64 accessions, with only one wild and cultivated accession, respectively. Hap3 was assigned to 44 accessions: 11 cultivated and 33 wild accessions. These distributions indicated that selection of *SiPLATZ12* likely occurred during domestication and breeding.

**Figure 3.**
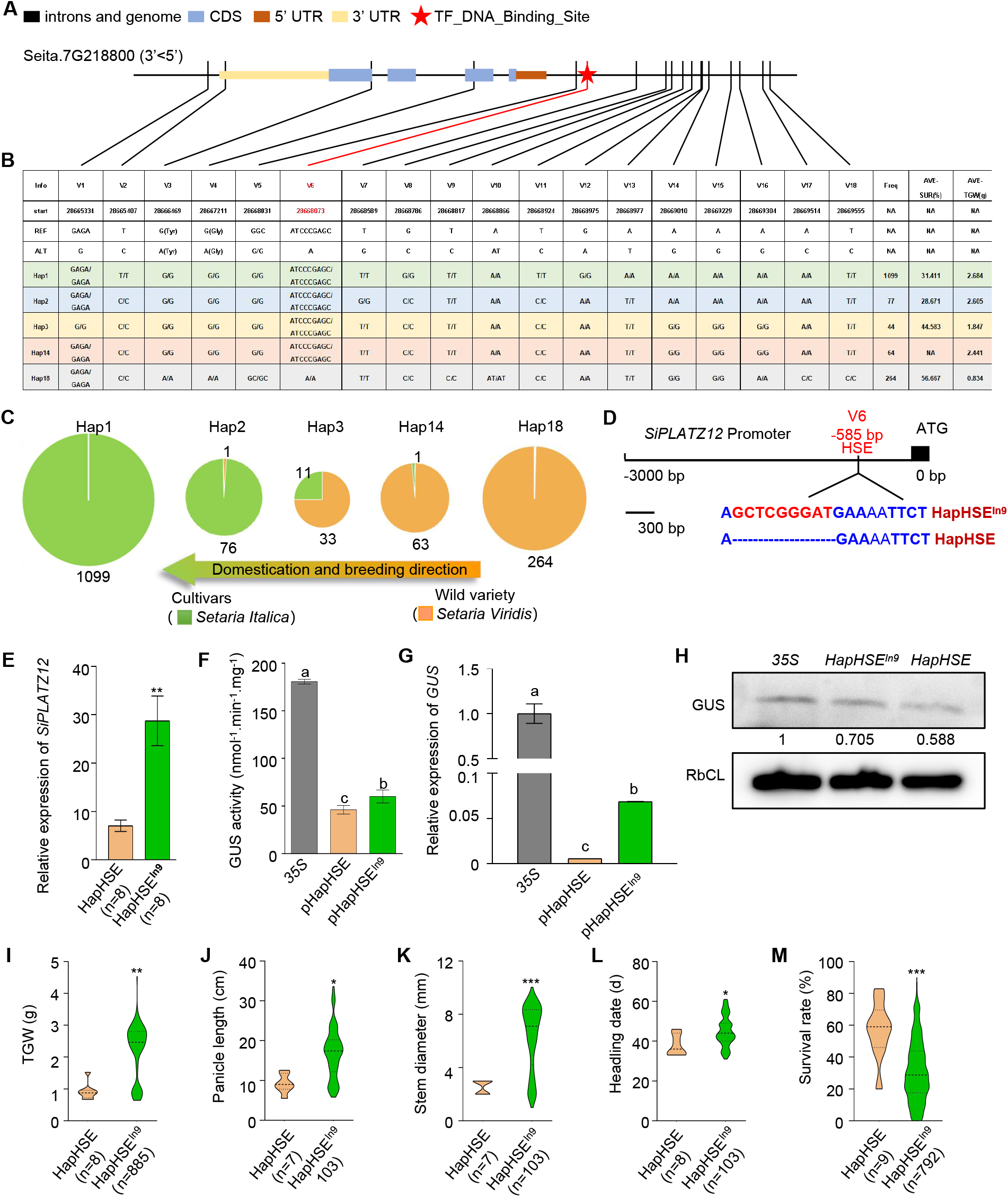
A 9-bp insertion in the *SiPLATZ12* promoter affects different expression of *SiPLATZ12*, yield traits and salt tolerance in millet. A Gene structure and distribution of DNA polymorphisms of *SiPLATZ12*. B Chromosome and corresponding positions of variant nucleotides in *SiPLATZ12*, five major haplotypes (Hap) of *SiPLATZ12*, and the corresponding average thousand-grain weight (AVE-TGW) under normal conditions and survival rate (AVE-SUR) under salt stress. REF and ALT represent reference sequence and alternative sequence, respectively. C Distribution of five major haplotypes among cultivated (*Setaria italica*) and wild (*Setaria viridis*) accessions. Numbers indicate frequencies (freq) of five major haplotypes. Arrow indicates domestication and breeding direction. D Two haplotypes caused by a 9-bp insertion (red letters) at -585 bp upstream of the translational start site of *SiPLATZ12*. HapHSE contains a correct heat stress response element (HSE) (blue letters), while HapHSE^In9^ contains a 9-bp insertion in HSE. E Relative expression levels of *SiPLATZ12* in every eight HapHSE and HapHSE^In9^ millet accessions, respectively. Three independent replicates were conducted. Values are given as the mean ± SD, \*\**P* < 0.01 by Student′s *t*-test. F GUS activities driven by HapHSE and HapHSE^In9^ promoters in tobacco leaves. *35S::GUS* was used as control. Data are presented as the mean ± SD of three replicates. *P* < 0.05 by one-way ANOVA. G, H Transcript (G) and protein (H) levels of *GUS* gene in tobacco leaves transformed with *pHapHSE::GUS* or *pHapHSE*^*In9*^*::GUS*, respectively. *35S::GUS* was used as control. Data are presented as the mean ± SD of three replicates. P < 0.05 by one-way ANOVA. I-M TGW (I), panicle length (J), stem diameter (K), heading date (L), SUR (M) of HapHSE and HapHSE^In9^ millet accessions. \**P* < 0.05; \*\**P* < 0.01; \*\*\**P* < 0.001by Student′s *t*-test.

To confirm the domestication and selection of *SiPLATZ12*, we analyzed the TGW among the five major haplotypes (Supplementary dataset 1). Hap1, Hap2, Hap3, and Hap14 millet accessions showed higher TGW (2.683, 2.605, 1.847, 2.441 g, respectively) than that of Hap18 accessions (0.834 g) (Fig 3B). The survival rate (SUR) of 801 millet accessions under salt stress were identified (Dataset EV1), and found lower SUR of Hap1, Hap2, and Hap3 (31.411%, 28.671%, 44.583%, respectively) than that of Hap18 accessions (56.667%) (Fig 3B). Obviously, the cultivated foxtail millet accessions have higher TGW but lower SUR than the wild ones, confirming the selection of *SiPLATZ12* during domestication and breeding.

To know whether and how *SiPLATZ12* is regulate in different millet varieties, we analyzed c*is*-element in the *SiPLATZ12* promoter and found a 9-bp insertion in a heat stress response element (HSE) (AGAANNTTCT) in Hap18 promoter resulted in Hap1, Hap2, Hap3, and Hap14 *SiPLATZ12* promoters (Fig 3B and D). Thus two haplotypes were generated at this site: HapHSE (AGAANNTTCT) and HapHSE^In9^ (A**GCTCGGGAT**GAANNTTCT). The distribution analysis showed that total of 1227 *S*.*italica* and 253 *S*.*viridis* accessions belong to HapHSE^In9^ type, while 413 *S*.*viridis* accessions belong to HapHSE type (Dataset EV1). This distribution of the two haplotypes indicated the selection of HSE^In9^ by breeders in domestication of *Setaria*.

To validate the regulation of *SiPLATZ12* by the 9-bp insertion in the HSE element, we detected the transcripts of *SiPLATZ12* in two haplotypic accessions, and found that *SiPLATZ12*^*HapHSEIn9*^ had higher transcripts than that of *SiPLATZ12*^*HapHSE*^ under normal condition but lower transcripts than that of *SiPLATZ12*^*HapHSE*^ under salt stress (Fig 3E, Appendix Fig S4A). These results indicated greater but lower salinity-inducible activity of the HapHSE^In9^ promoter than the HapHSE promoter. Consistently, the HapHSE^In9^ promoter drove higher GUS activity than that of the HapHSE promoter in tobacco (*Nicotiana tabacum*) epidermal cells (Fig 3F). And higher GUS transcript and protein levels driven by the HapHSE^In9^ promoter were detected in tobacco epidermal cells than those driven by the HapHSE promoter (Fig 3G and H). Importantly, *SiPLATZ12*^HapHSEIn9^ was positively associated with TGW, panicle length, main stem diameter, and heading time, while *SiPLATZ12*^HapHSE^ was positively correlated with SUR (Fig 3I-M, Dataset EV1). The changes in TGW and SUR of some represent HapHSE^In9^ and HapHSE accessions, and expression levels of *SiPLATZ12* were illustrated in Appendix Fig S4B-F. These results indicated that variation in *cis*-regulation of *SiPLATZ12* contribute to its differential expression, and thus diverse yield traits and salt tolerance in millet.

### SiPLATZ12 regulates the expression of seed development and salt tolerance related genes

To explore the mechanism of *SiPLATZ12* in regulating seed development and salt stress response in foxtail millet, the expression of the related genes was detected. The transcript levels of *IKU1, IKU2, MINI3, SHB1, KLU, FLS2*, and *CKI1* were obviously up-regulated by *SiPLATZ12* overexpression in foxtail millet (Fig 4A). In contrast, transcript levels of most *SiNHX, SiSOS*, and *SiCBL* genes were significantly reduced by *SiPLATZ12* overexpression in foxtail millet under both normal and salt-stress conditions (Fig 4B and C). Similar expression patterns for *AtNHX, AtSOS*, and *AtCBL* genes were obtained in *SiPLATZ12* transgenic *Arabidopsis* seedlings when compared with Col-0, except for *AtCBL9* (Appendix Fig S5 and 6). Therefore, SiPLATZ12 increases the expression of genes controlling seed size, but inhibits that of *NHX, SOS*, and *CBL* genes in both foxtail millet and *Arabidopsis*.

**Figure 4.**
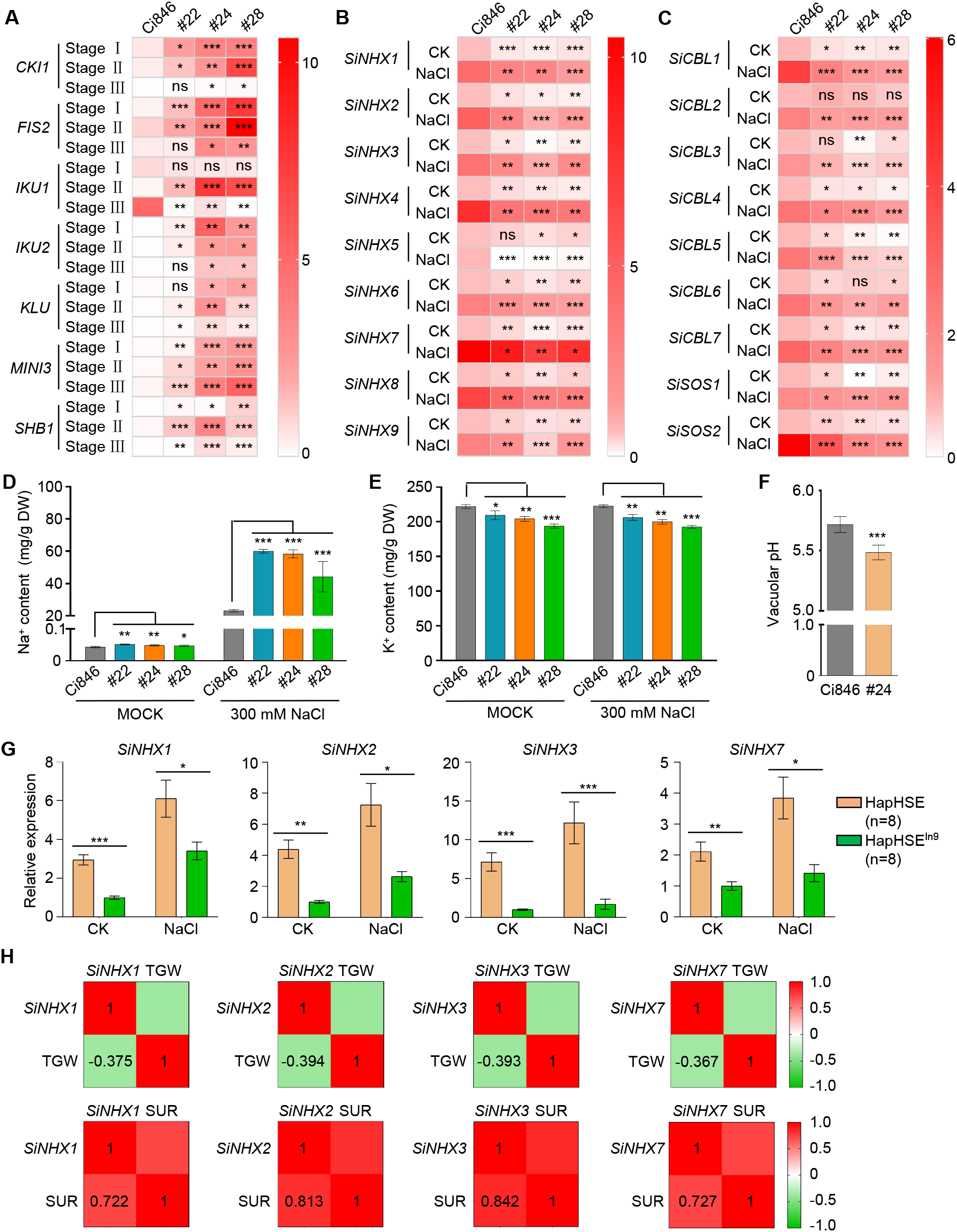
SiPLATZ12 regulates the expression of seed development and salt tolerance related genes. A Absolute expression levels of genes related to seed development in panicles at 3 day (Stage I), 10 day (stage II), and 20 day (stage III) after heading of ‘Ci846’ or *SiPLATZ12* overexpression foxtail millet. *SiACTIN7* and *18S rRNA* were used as internal controls. Color scale for log2 FC (fold change) values is shown at the right. Experiments were performed in biological triplicate. \**P* < 0.1; \*\**P* < 0.01; \*\*\**P* < 0.001 by Student′s *t*-test. B, C *SiNHXs, SiSOSs* and *SiCBLs* in 7-day old ‘Ci846’ or *SiPLATZ12* overexpression foxtail millet treated with or without 200 mM NaCl for 24 h. *SiACTIN7* and *18S rRNA* were used as internal controls. Color scale for log2 FC values is shown at the right. Experiments were performed in biological triplicate. \**P* < 0.1; \*\**P* < 0.01; \*\*\**P* < 0.001 by Student′s *t*-test. D, E Na^+^ and K^+^ contents of ‘Ci846’ and *SiPLATZ12* overexpression millet seedlings treated with or without 300 mM NaCl for 6 h. Data are presented as the mean ± SD of three replicates. \**P* < 0.05, \*\**P* < 0.01, \*\*\**P* < 0.001 by Student′s *t*-test. F Vacuolar pH in roots of 4-d etiolated seedlings of ‘Ci846’ and *SiPLATZ12* overexpression foxtail millet (#24), and measured by BCECF-AM. Data are presented as the mean ± SD; n > 10. \*\*\**P* < 0.001 by Student′s *t*-test. G Relative expression of *SiNHX1, SiNHX2, SiNHX3 and SiNHX7* in HapHSE and HapHSE^In9^ millet accessions under normal and salt stress conditions. Data are presented as the mean ± SD of three replicates. \**P* < 0.05, \*\**P* < 0.01, \*\*\**P* < 0.001 by Student′s *t*-test. H Correlations of TGW and SUR with relative expression levels of *SiNHX1, SiNHX2, SiNHX3* and *SiNHX7* under normal and salinity conditions.

Since NHXs act in cell expansion by regulating Na^+^, K^+^ and vacuolar pH homeostasis (Bassil *et al*., 2011a; Bassil *et al*., 2011b), we measured the Na^+^, K^+^ and pH changes in ‘Ci846’ and *SiPLATZ12*-overexpressing foxtail millet seedlings. We observed higher Na^+^ contents in the transgenic foxtail seedlings than in ‘Ci846’ under both normal and salinity conditions (Fig 4D). In contrast, K^+^ contents in *SiPLATZ12*-overexpressing foxtail millet were lower than those in ‘Ci846’ under both normal and salinity conditions (Fig 4E). Although no significant difference in vacuolar pH between root tip cells of ‘Ci846’ and *SiPLATZ12*-overexpressing seedlings, vacuolar pH in cells of the mature root zone was nearly 0.25 units lower in SiPLATZ12-overexpressing seedlings compared with ‘Ci846’ seedlings (Fig 4F, Appendix Fig S7). These data suggested that SiPLATZ12 disturb Na^+^, K^+^ and vacuolar pH homeostasis through inhibiting the expression of *NHX* and related genes in foxtail millet.

To establish the relationship of *SiNHX1, SiNHX2, SiNHX3*, and *SiNHX7* with seed size and salt tolerance in foxtail millet, we detected the expression of them in HapHSE and HapHSE^In9^ millet accessions, respectively. The results exhibited that transcript levels of *SiNHX1, SiNHX2, SiNHX3*, and *SiNHX7* in seedlings of HapHSE millet accessions was significantly higher than that in HapHSE^In9^ millet accessions under both normal and salinity conditions (Fig 4G). Then, we performed correlation analysis and found that the TGW of these accessions showed negative and weak correlations (0.3<R^2^<0.5), while the SUR of these accessions showed positive and strong correlations (R^2^>0.7) with the expression levels of the four genes (Fig 4H), indicating negative and partial contribution of *SiNHX1, SiNHX2, SiNHX3* and *SiNHX7* to TGW while positive and main contribution of them to SUR in millet.

### SiPLATZ12 binds directly to the promoters of *SiNHX2, SiCBL4* and *SiCBL7*

To test whether SiPLATZ12 acts as a direct transcription regulator of genes mentioned above, we verified the direct binding of SiPLATZ12 to the promoters of *SiNHX2, SiCBL4*, and *SiCBL7* by yeast one hybrid (Y1H) experiment (Fig 5A). After the confirmation of the nuclear localization of SiPLATZ12 in foxtail millet hairy-roots and transgenic *Arabidopsis* roots (Appendix Fig S8A and B), we performed chromatin immunoprecipitation (ChIP)-qPCR analysis using the SiPLATZ12-GFP transgenic hairy roots of foxtail millet using specific primers for cloning A/T rich DNA fragments within the promoters of *SiNHX2, SiCBL4*, and *SiCBL7* (Table S1, Appendix Fig S9A). The results showed that greater enrichment of P4 and P5 DNA fragments within the *SiNHX2* promoter, P7 and P9 DNA fragments within the *SiCBL4* promoter, and P10 and P12 within the *SiCBL7* promoter were obtained in pPLATZ12::PLATZ12-GFP transgenic samples than in pPLATZ12::GFP transgenic samples, while other indicated fragments showed no difference between them (Fig 5B-D), indicating the binding of SiPLATZ12 to *SiNHX2, SiCBL4*, and *SiCBL7* promoters *in vivo*. The results of microscale thermophoresis (MST) assays between SiPLATZ12 and P5, P9, and P10 DNA fragments within *SiNHX2, SiCBL4*, and *SiCBL7* promoters confirmed the binding of SiPLATZ12 to *SiNHX2, SiCBL4*, and *SiCBL7* promoters *in vitro* (Fig 5E, Appendix Fig S9B). Finally, we performed electrophoretic mobility shift assays (EMSAs) and found that SiPLATZ12-His shifted all of the labeled *SiNHX2, SiCBL4*, and *SiCBL7* probes (Fig 65F). These shifted probes were gradually weakened by increasing concentrations of unlabeled *SiNHX2, SiCBL4*, and *SiCBL7* DNA fragments until they were no longer detectable. Meanwhile, no shifted bands were observed when *SiNHX2, SiCBL4*, or *SiCBL7* probes were mutated to C/G DNA probes. These data proved direct binding of SiPLATZ12 to some A/T rich sequences in the promoters of *SiNHX2, SiCBL4*, and *SiCBL7 in vitro*.

**Figure 5.**
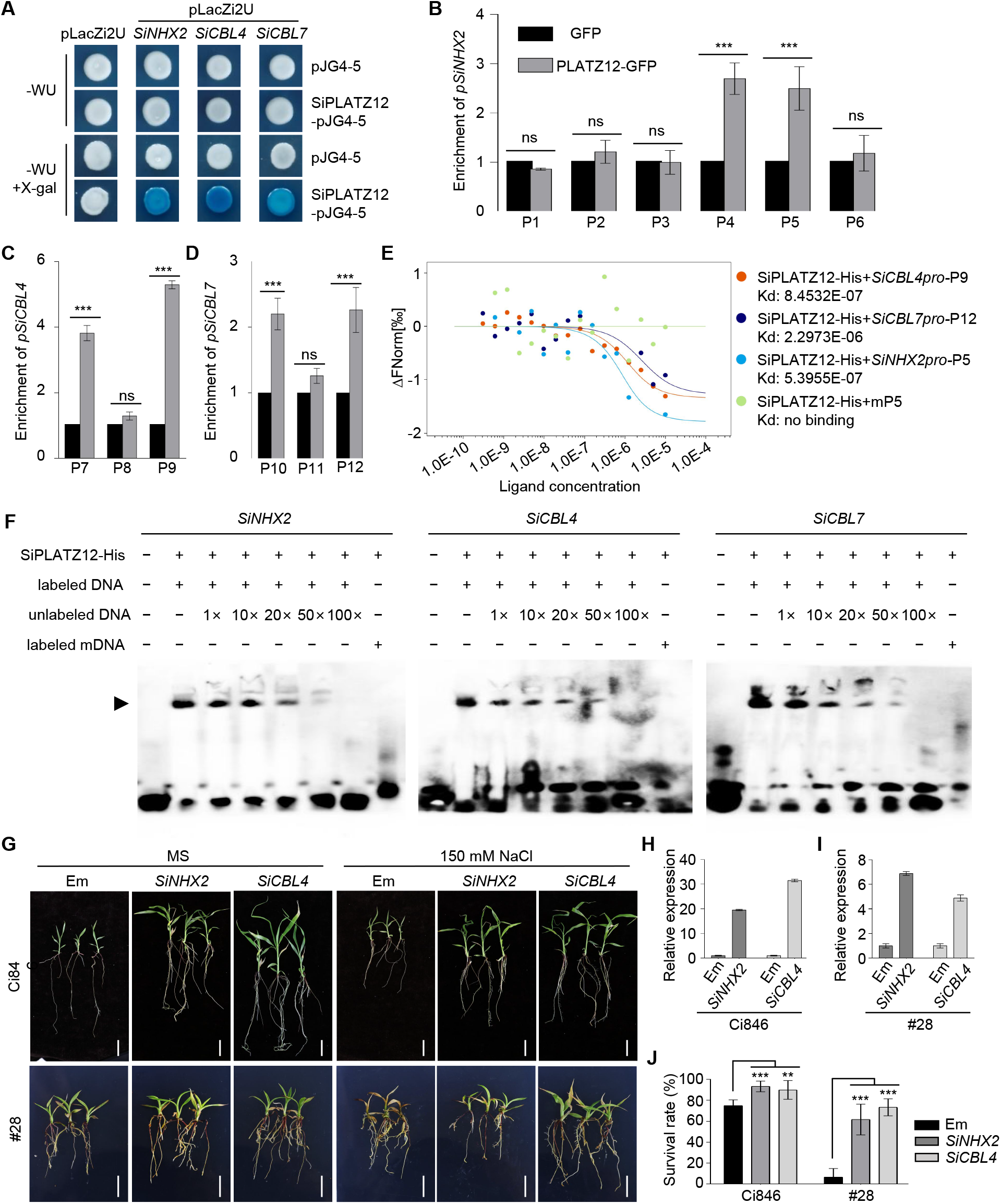
SiPLATZ12 binds directly the promoters of *SiNHX2, SiCBL4*, and *SiCBL7* genes. A Yeast one-hybrid assays show the binding of SiPLATZ12 to *SiNHHX2, SiCBL4*, and *SiCBL7* promoters. WU, Synthetic Dropout/-Trp-Ura. B-D ChIP-qPCR analysis determine the SiPLATZ12 binding regions in the *SiNHHX2, SiCBL4* and *SiCBL7* promoters. Data are presented as the mean ± SD of three replicates. ns, not significant; \*\*\**P* < 0.001 by Student′s *t*-test. E MST assays verify the direct binding of SiPLATZ12 to the selected selected A/T sequences in *SiNHHX2, SiCBL4*, and *SiCBL7* promoters. A mutant DNA fragment (mP5) was used as control. F EMSAs verify the binding of SiPLATZ12 to the selected A/T sequences in *SiNHHX2, SiCBL4* and *SiCBL7* promoters. DNA sequences and the corresponding mutant sequence used in this experiment were listed in Table S2. Arrow indicates shifted band of DNA by SiPLATZ12. The plus (+) or minus (-) denote the presence or absence of SiPLATZ12 or DNA in each sample. G Photographs of ‘Ci846’ and ‘#28’ *SiPLATZ12* overexpression seedlings transformed with empty vector (EM), *35S::SiNHX2* (*SiNHX2*) or *35S::SiCBL4* (*SiCBL4*) and cultured on MS medium with or without 150 mM NaCl for 7 days. Scale bars = 1 cm. Data represents the mean ± SD of three replicates. \*\**P* < 0.01, \*\*\**P* < 0.001 by Student′s *t*-test. H, I Expression levels of *SiNHX2* and *SiCBL4* in transgenic ‘Ci846’ (H) and ‘#28’ (I) seedlings. *SiACTIN7* and *18S rRNA* were used as the internal control. Data represents the mean ± SD of three replicates. **P < 0.01, ***P < 0.001 by Student′s *t*-test. J Survival rates of ‘Ci846’ and ‘#28’ *SiPLATZ12* overexpression seedlings transformed with EM, *SiNHX2* or *SiCBL4* and cultured on MS medium with or without 150 mM NaCl for 7 day. At least 30 seedlings were used. Data represents the mean ± SD of three replicates. \*\**P* < 0.01, \*\*\**P* < 0.001 by Student′s *t*-test.

To determine the genetic relationship of *SiPLATZ12* and *SiNHX2* and *SiCBL4, SiNHX2* and *SiCBL4* were overexpressed in hairy-roots of foxtail millet overexpressing *SiPLATZ12* (#28) and ‘Ci846’. As expected, all ‘Ci846’ and #28 seedlings with *35S::SiNHX2* and *35S*::*SiCBL4* transgenic hairy roots grew better than with empty vector transgenic hairy roots (Fig 5G-I) under normal conditions. Meanwhile, SURs of seedlings with *SiNHX2* and *SiCBL4* transgenic hairy roots were 55% and 66.7% higher than that with the empty vector transgenic hairy roots under salt stress (Fig. 5G and J). These observations support that *SiPLATZ12* functions upstream of *SiNHX2* and *SiCBL4*.

## Discussion

Seed size is a major factor controlling grain yield, and is strongly affected by various genetic factors, such as the IKU pathway, the ubiquitin-proteasome pathway, G-protein signaling, the mitogen-activated protein kinase signaling pathway, phytohormones, and transcriptional regulatory factors in Arabidopsis (Kesavan *et al*., 2013; Li and Li, 2016). We revealed that *SiPLATZ12* play important roles in regulating seed size in foxtail millet (Fig 1A-D; Fig 3B and I) by upregulating the expression of genes involved in the IKU pathway, including *IKU1, IKU2, MINI3*, and *SHB1* (Fig 4A). Loss of function of *IKU1, IKU2, MINI3*, and *SHB1* genes reduces seed size due to precocious cellularization of the endosperm (Luo *et al*., 2005; Wang *et al*., 2010; Zhou *et al*., 2009). Therefore, SiPLATZ12 is a novel activator of genes in the IKU pathway to influence endosperm growth or grain filling. Moreover, *KLU* encodes the cytochrome P450 CYP78A5 and positively regulates seed size by promoting cell proliferation in maternal integuments (Adamski *et al*., 2009). The up-regulated expression of *KLU* by *SiPLATZ12* overexpression in foxtail millet (Fig 4A) indicating a role of *SiPLATZ12* in maternal control of seed size at least through activating *KLU* expression. Besides, FL3 and GL6 are required for tRNA and 5S rRNA transcription through interaction with RNA Polymerase III (Kim *et al*., 2018; Li *et al*., 2017; Wang *et al*., 2019; Wang *et al*., 2018). Consistent with the binding sequences of FL3 and GL6, SiPLATZ12 bound A/T rich sequences in the promoters of target genes (Fig 5B-F), indicating possible similar mechanisms of SiPLATZ12 with FL3 and GL6 in regulating seed size. Thus, SiPLATZ12 controls seed length and width by regulating the expression of multiple genes involved in several signaling pathways.

Panicle size also strongly affects grain yield. SiMADS34, E-class MADS-box TF, positively regulates panicle length but negatively regulates panicle width and grain yield in *S*.*italica* by upregulating Seita.6G205500, a homology of rice SPL14 (Yang *et al*., 2021). A recessive mutant of *Loose Panicle 1* (*LP1*), which encodes a group I WRKY TF, shows pleiotropic phenotypes, such as a lax primary branching pattern, aberrant branch morphology, semidwarfism, long and wide panicles, and big seeds (Xiang *et al*., 2017). Similar to LP1 but different from SiMADS34, SiPLATZ12 positively regulates panicle size and grain number in foxtail millet (Fig 1E-H), reflecting the complexity of panicle and grain number development.

Moreover, plant height and stem diameter are usually thought to be associated with high grain yield by changing lodging resistance of most crops. Breeding of the varieties carrying the dwarfing genes (*Rht*) is the main direction to reduce the risk of lodging and increase grain yield (Evans, 1998; Rebetzke *et al*., 2011; Tang *et al*., 2021). The diameter of the stem correlates with resistance to lodging in rice and wheat (Ageeva *et al*., 2020; Kashiwagi *et al*., 2008). The loci with high phenotypic effects on lodging tolerance are colocalized with loci responsible for plant height, stem diameter and stem strength (Ageeva *et al*., 2020). Due to the reduced plant height and improved stem diameter of *SiPLATZ12* overexpression millet (Fig 1J-M, Fig 3K; Dataset EV1), SiPLATZ12 may be a novel loci controlling lodging resistance.

Besides, heading time is an important developmental transition in plants leading to successful sexual reproduction and is determined by multiple genes and environmental factors, such as day-length and temperature (Freytes *et al*., 2021). Several genetic factors controlling the heading date of foxtail millet have been identified. *Heading date 1* (*HD1*), homolog gene of *CO*, has been identified as a candidate of a quantitative trait loci (QTL) for heading date using genome-wide association studies (Jia *et a*l., 2013), which is under strong selection during domestication (Liu *et a*l., 2015). *Roc4* was identified by QTL sequence (Yoshitsu *et al*., 2017). Roc4 promotes flowering time under long days in rice (Wei *et al*., 2016). The delayed heading of *SiPLATZ12* transgenic foxtail millet and the association of heading date with HapHSE^In9^ indicate a novel genetic factor in controlling heading date and the domestication of SiPLATZ12^HapHSEIn9^ in foxtail millet (Fig 1JL, N, Fig 3L; Dataset EV1).

In addition, SiPLATZ12 negatively regulates millet salt tolerance. The salt tolerance of plants has been proved to be largely associated with cytoplasm Na^+^ homeostasis which is maintained by membrane localized NHXs and their regulatory proteins, such as SOS2 and CBLs (Halfter *et al*., 2000; Liu and Zhu, 1998; Quintero *et al*., 2011; Zhou *et al*., 2022). We found that SiPLATZ12 repressed the expression of most of *NHX, SOS* and *CBL* genes, and directly targeted *SiNHX2, SiCBL4*, and *SiCBL7* genes (Fig 4B and C; Fig 5A-F; Appendix Fig S9). *SiNHX2* and *SiCBL4* overexpression could rescue the salt sensitivity of *SiPLATZ12* overexpression foxtail millet (Fig 5G-J). Therefore, *SiPLATZ12* overexpression might lead to lower Na^+^/H^+^ antiporter activities of SOS1 and NHXs, which in turn leads to the increased Na^+^ accumulation in *SiPLATZ12-*overexpressing foxtail millet (Fig 4D). Simultaneously, NHXs participate in vacuolar K^+^ and pH homeostasis (Bassil *et al*., 2011a; Bassil *et al*., 2011b). NHX5 and NHX6-mediated pH increasing of endomembrane compartments influences the sorting of transmembrane proteins (Bassil *et al*., 2011a). Thus, NHX operations are essential for many processes, including stomatal regulation (Barragán *et al*., 2012), flower development (Bassil *et al*., 2011b; Yoshida *et al*., 2005), plant reproduction (Bassil *et al*., 2012; Bassil *et al*., 2011b), and seed development (Reguera *et al*., 2015). Consistently, Na^+^, K^+^ and vacuolar pH homeostasis were disturbed in foxtail millet overexpressing *SiPLATZ12* (Fig 4D-F), which might be the reason behind the multiple changed yield traits under normal conditions (Fig 1). Furthermore, TGW showed negative correlation while salt tolerance showed positive correlation with the expression levels of *SiNHX1, SiNHX2, SiNHX3*, and *SiNHX7* (Fig 4G and H). Therefore, SiPLATZ12 regulates multiple yield traits and salt tolerance partially by suppressing the expression of *NHX, SOS*, and *CBL* genes.

Tradeoffs among agronomic traits limit the crop yield, such as penalties of yield by plant immunity, grain quality by yield and negative correlations among plant architecture components (Nelson *et al*., 2018; Wang *et al*., 2021; Song *et al*., 2022). Many of these tradeoffs are caused by gene pleiotropy. Our results provide strong evidence for the *SiPLATZ12* pleiotropy (Fig 1, 2 and 3, Appendix Fig S3). Two tradeoffs between TGW and salt tolerance, and grain number per panicle and seed size were caused by SiPLATZ12. Changes on *cis*-regulatory regions can regulate quantitative traits at different levels (Rodriguez-Leal *et al*., 2017; Hendelman *et al*., 2021; Liu *et al*., 2021). The yield and salt tolerance are determined in different conditions, while panicle size and seed size are determined in different developmental stages. Therefore, modifying *cis*-regulatory regions of *SiPLATZ12* may overcome the drag between yield and salt tolerance as well as grain number per panicle and seed size to outperform current elite natural alleles. In fact, a 9-bp insertion in the *SiPLATZ12* promoter disrupted the HSE element and resulted in higher expression of *SiPLATZ12*^*HapHSEIn9*^ than *SiPLATZ12*^*HapHSE*^ under normal condition (Fig S4F), supporting the efficiency of overcoming the tradeoffs between different yield traits by modifying *cis*-regulatory regions of *SiPLATZ12*. Therefore, identifying and modifying the *cis*-elements responding to salt stress and heading date may reduce the pleiotropic effect of *SiPLATZ12* and generate a ‘ideal’ phenotype variation in foxtail millet.

In summary, our study reveals the genetic mechanisms by which SiPLATZ12 promotes multiple elite yield traits at the expense of salt tolerance in foxtail millet (Fig 6). The expression levels of *SiPLATZ12* in HapHSE^In9^ millet accessions (landrace and cultivated varieties) are increased by a 9-bp insertion in the promoter of *SiPLATZ12*. On one hand, SiPLATZ12 then activates the expression of genes in the IKU pathway, including *IKU1, IKU2, MINI3*, and *SHB1*, and the maternal control genes, such as *KLU*, to increase seed size. On the other hand, SiPLATZ12 inhibits the expression of *NH*X, *SOS*, and *CBL* genes, at least directly targets to *SiNH*X2, *SiCBL4* and *SiCBL7*, to increase TGW, panicle size and stem diameter, but inhibit plant height, heading date and salt tolerance of foxtail millet. Our results provide novel insights into the genetic basis of the trade-off between yield traits and salt tolerance in millet. However, further elucidating the regulatory relationships of SiPLATZ12 with other regulators controlling yield traits and salt tolerance modifying the *cis*-element of *SiPLATZ12* will contribute well to breeding of new elite rice cultivars with “ideal” plant architecture to improve grain yield of foxtail millet.

**Figure 6.**
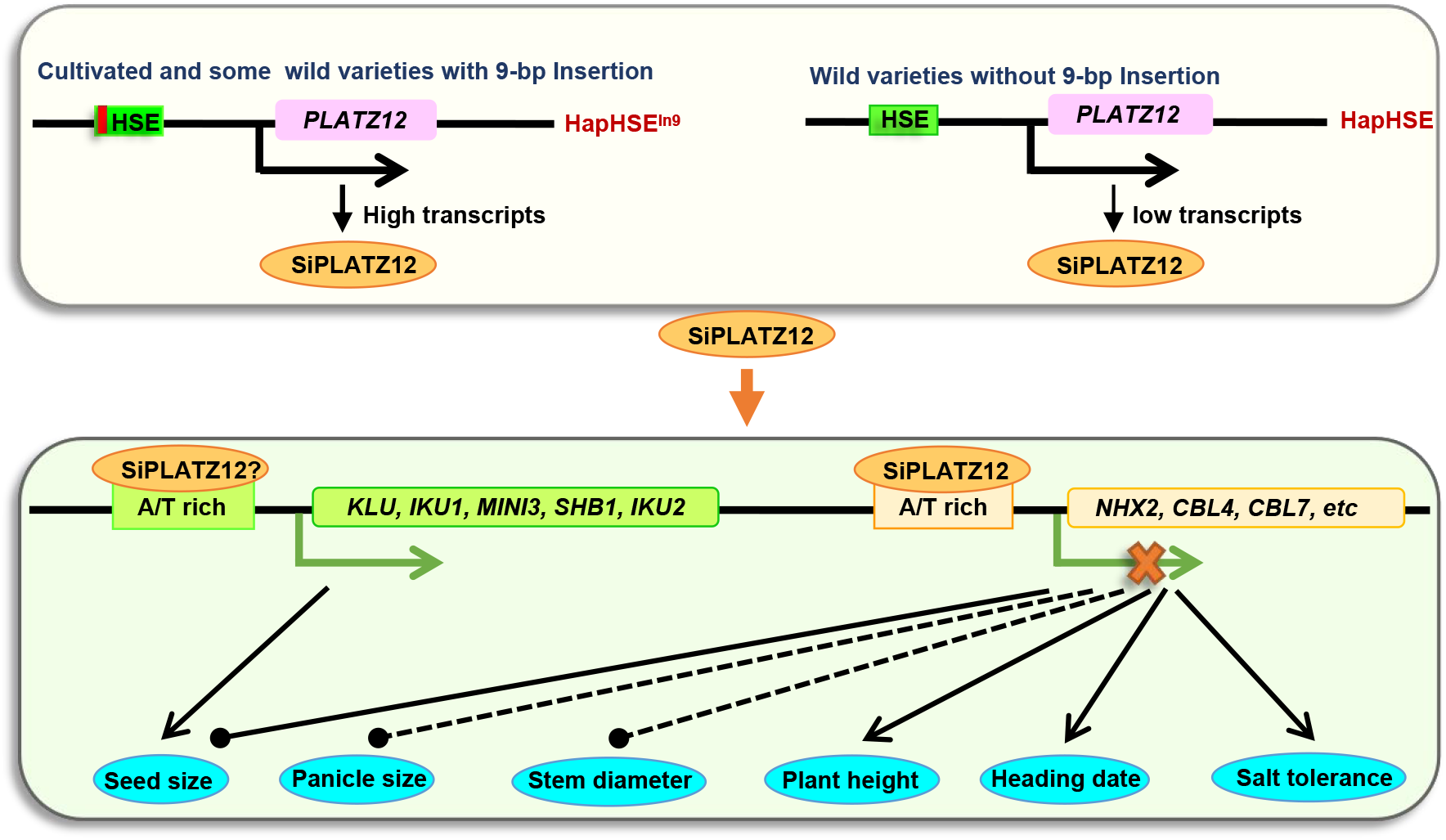
A model for SiPLATZ12-regulated yield traits and salt tolerance in millet. A 9-bp insertion in the *SiPLATZ12* promoter leads to the increased expression of *SiPLATZ12* in HapHSE^In9^ millet accessions (landrace and cultivated varieties) compared to HapHSE (wild) accessions. On one hand, SiPLATZ12 directly targets and inhibits the expression of *NHX, SOS*, and *CBL* genes, leading to increased panicle size and TGW, but reduced plant growth and salt tolerance in foxtail millet. On the other hand, SiPLATZ12 activates the expression of genes involved in the IKU pathway, including *IKU1, IKU2, MINI3*, and *SHB1*, and in the maternal control, such as *KLU*, to increase seed size.

## Materials and Methods

### Plant materials and growth conditions

Foxtail millet (*S. italica*) cultivars ‘Yugu1’ and ‘Ci846’, and *Arabidopsis* Col-0 were used for gene expression and function analyses. Foxtail millet seeds were sterilized with sodium hypochlorite, washed three times and germinated at 28°C for 3 d. For inducible expression analysis, 10-day-old seedlings of ‘Yugu1’ were treated with Hoagland solution containing 200 mM NaCl, 50 mM NaHCO_3_, 100 μM ABA, and 20% (w/v) polyethylene glycol 6000 (PEG6000) or at 4°C or 42°C for 0, 3, and 24 h. After treatment, whole seedlings were collected and frozen quickly in liquid nitrogen for RNA extraction. For tissue expression analysis, roots, stems and leaves from 10-day-old seedlings, panicles from 2-month-old seedlings and seeds of ‘Yugu1’ were collected and frozen quickly in liquid nitrogen. For phenotypic analysis, seeds were germinated on filter paper wetted with water or salt solution for the times indicated. Alternatively, the 3-day-old uniformly developed seedlings were cultured for 10 d under 16/8 h light/dark period and 28 ± 1°C or in the field during summer for the times indicated. Transgenic hairy roots were prepared as described previously (Zhang *et al*., 2021). Culture *Arabidopsis* materials were described previously (Yu *et al*., 2019). Phenotypes were photographed, and root or shoot length and fresh weight were measured.

For salt tolerance haplotype analysis, 801 germplasms, including wild, landbrace and cultivated accessions, were used (Supplementary dataset1). Four-day-old seedlings were transferred to pots with matrix (nutrient soil: vermiculite=1:1) for 7 d. Healthy developed seedlings were then saturated in 300 mM NaCl solution for three consecutive times (once every 3 days) and grown for an additional 10 d. Surviving seedlings were counted, and survival rates were calculated. Three independent experiments were carried out.

### Identification of *SiPLATZ, SiNHX, SiSOS1, SiSOS2* and *SiCBL* genes in foxtail millet

*PLATZ* and *NHX* family members and *SOS2* from *Arabidopsis* were used to BLAST the foxtail millet genome in Phytozome V12 (https://phytozome.jgi.doe.gov) to identify *SiPLATZ, SiNHX* and *SiSOS2* genes. *SiSOS1* (Yan *et al*., 2021) and S*iCBL* genes (Zhao *et al*., 2013) has been identified. Isoelectric point and molecular weight were predicted using the ExPASy Proteomics Server (http://expasy.org/). An evolutionary tree was generated using MEGA 6 software and a phylogenetic evolutionary tree was constructed using the neighbor-joining analysis and bootstrap method, with 1000 replicates.

### RNA extraction and quantitative RT-PCR (RT-qPCR)

Total RNA was extracted using RNAiso Plus (Takara, Ohtsu, Japan). As described previously (Yu *et al*., 2019), reverse transcriptions were performed using PrimeScriptTM RT reagent kit (TaKaRa, Ohtsu, Japan). RT-qPCR was performed using ChamQ Universal SYBR qPCR Master Mix (Vazyme, Biotech Co., Ltd) in a three-step program on a CFX96TM Real-Time PCR Detection System (Bio-Rad, Hercules, CA, USA). 18S ribosomal RNA and *SiACT7* (Li *et al*., 2016) were used as internal controls. Three biological replicates were performed. Primers are shown in Table S1.

### Generation of transgenic foxtail millet and *Arabidopsis*

Transgenic foxtail millet (cultivar ‘Ci846’) plants were generated by introducing the gene-coding region of *SiPLATZ12* under control of the *35S* promoter via callus-based gene transformation using *Agrobacterium tumefaciens* strain GV3101. Transformants were selected on half-strength MS medium supplemented with 50 mg/L kanamycin and planted to soil for harvesting seeds. Transgenic foxtail millet seedlings were confirmed by RT-qPCR using primers listed in Table S1.

Transgenic *Arabidopsis* containing *35S::SiPLATZ12* and *35S*::*SiPLATZ12-GFP* was generated simultaneously using the floral-dipping method. T_3_ transgenic *Arabidopsis* plants were selected on half-strength MS medium supplemented with 50 mg/L kanamycin and confirmed by RT-qPCR using primers listed in Table S1.

### Haplotype analysis

We collected the available data for phenotypes related to yield traits for 916 accessions in our previous study (Jia *et al*., 2013) The genotype data for those accessions were obtained by high-depth resequencing (unpublished). The DNA sequences, comprising promoter (3 kb upstream of ATG), gene body (2360 bp between 28665353 to 28667712 in chromosome 7, reverse), and 2 kb downstream of *SiPLATZ12*, were extracted. The haplotype analysis was performed using in-house python and R scripts.

### Fluorometric GUS assays

*pHapHSE*^*In9*^ and *pHapHSE* representing the 869 bp intergenetic region upstream of ATG of *SiPLATZ12* from *HapHSE*^*In9*^ and *HapHSE* millet accessions, and was fused to the *pBI121* vector to drive *GUS* expression. Then *pHapHSE*^*In9*^*::GUS* and *pHapHSE::GUS* constructs were transiently transformed into tobacco (*N. benthamiana*) leaves. GUS enzymatic activities in tobacco leaves were measured on an F-4500 fluorescence spectrofluorometer (Hitachi, Tokyo, Japan) using 4-methylumbelliferyl-β-d-glucuronide as a substrate. Standard curves were prepared using 4-methylumbelliferone. Mean GUS activities were obtained from three independent measurements, and each assay was repeated three times.

### Immunoblotting analysis

Root samples (0.5 g) of 7-day-old transgenic foxtail millet seedlings harboring *pHapHSE*^*In9*^*::GUS* and *pHapHSE::GUS* obtained by via *Agrobacterium tumefaciens* transformation were ground in liquid nitrogen and mixed with 500 μl of non-denatured protein extract [Tris–HCl (pH 7.5), 25 mM; MgCl_2_, 10 mM; NaCl, 10 mM; DL-Dithiothreitol (DTT), 5 mM; phenylmethylsulfonyl fluoride (PMSF), 4 mM; ATP, 10 mM]. The crude extract was placed on ice for 30 min and then centrifuged at 17 000 *g* for 15 min at 4°CC. The supernatant was centrifuged again, and the supernatant was collected. Anti-GUS antibody was used to examine the protein level of GUS.

### Y1H assay

*SiNHX2, SiCBL4*, and *SiCBL7* promoters were amplified and cloned into pLacZi2μ, which contains the *LacZ* reporter gene, to generate *pSiNHX2::LacZ, pSiCBL4::LacZ* and *pSiCBL7::LacZ*, respectively. The cDNA sequence (CDS) of *SiPLATZ12* was cloned into the pJG4-5 vector containing the GAL4 transcriptional activation domain (GAD) to generate *PLATZ12-pJG4-5. PLATZ12-pJG4-5* and *pSiNHX2::LacZ, pSiCBL4::LacZ* or *pSiCBL7::LacZ* were co-transformed into yeast strain EGY48 according to the Yeast Protocols Handbook (Clontech). Transformed yeast (*Saccharomyces cerevisiae*) cells were plated onto selective medium without Ura or Trp, and positive clones were cultured on medium without Ura or Trp containing galactose (20%), raffinose (20%), BU-salt (50 ml: 1.95 g Na_2_HPO_4_, 1.855 g NaH_2_PO_4_·2H_2_O), and 5-bromo-4-chloro-3-indolyl-β-D-galactopyranoside (X-Gal) for development of blue colour. Yeast cells co-transformed with pJG4-5 and *pSiNHX2::LacZ, pSiCBL4::LacZ* or *pSiCBL7::LacZ* were used as negative controls.

### Subcellular localization analysis

To determine the subcellular localization of SiPLATZ12, transgenic foxtail millet hairy roots and *Arabidopsis* harboring *35S::SiPLATZ12-GFP* were generated. GFP fluorescence was then observed using a LSM880 high-resolution laser confocal microscope (Zeiss, Germany). DAPI (4′,6-diamidino-2-phenylindole) was used to stain the nucleus.

### ChIP-qPCR assay

ChIP assays were performed as described previously (Liu *et al*., 2020). Seven-day-old *35S::SiPLATZ12-GFP* transgenic seedlings and GFP-specific monoclonal antibody (Beyotime Biotechnology, China) were used for ChIP. ChIP DNA products were analyzed by qPCR with primers designed to amplify the indicated DNA fragments within the *SiNHX2, SiCBL4* and *SiCBL7* promoter, respectively.

### MST assay

MST assay was performed according to the method established by Wienken et al. (2010). SiPLATZ12-His proteins were induced in *Escherichia coli* Rosetta cells and purified using a His-Tagged Protein Purification Kit (CWBIO, Beijing, China) as described previously (Liu et al., 2020). Binding reaction of recombinant SiPLATZ12-His protein to different probes was measured by microscale thermophoresis (MST) in a Monolith NT.115 (Nano Temper Technologies) instrument. The SiPLATZ12-His labeled with Cy5 were diluted to 40 nM by a buffer containing 50 mM Tris-HCl (pH 7.4) and 0.05% (v/v) Tween 20. A range of concentrations of different probes in the assay buffer [50 mM Tris-HCl (pH 7.8), 150 mM NaCl, 10 mM MgCl_2_, 0.05% Tween 20] was incubated with labeled protein (1:1, v/v) for 10 min. The sample was loaded into the Monolith NT.115 standard capillaries and measured with 40% MST power. The KD Fit function of the Nano Temper Analysis Software was used to fit the curve and calculate the value of the dissociation constant (Kd).

### EMSA

Three indicated DNA fragments within the *SiNHX2, SiCBL4* and *SiCBL7* promoters were respectively amplified using biotin-labeled primers (Table S1) synthesized by Sangon Biotech (Shanghai China) and purified using a PCR purification kit (Qiagen). EMSA was conducted using a LightShift Chemilumniescent EMSA Kit (Thermo Fisher Scientific, Pierce, USA) as described previously (Liu *et al*., 2020). Unlabeled DNA fragments of the same sequences were used as competitors. Migration of biotin-labeled probes was detected using enhanced chemiluminescence substrate (Thermo Scientific) on a ChemDoc XRS system (Bio-Rad).

### Na^+^ and K^+^ content measurement

To determine Na^+^ and K^+^ contents, one-week-old seedlings were treated with 200 mM NaCl for 12 h before harvesting, drying for 48 h at 80°CC and then grinding to powder. Tissue powder (100 mg) was digested in concentrated nitric acid, hydrochloric acid and hydrogen peroxide for 30 min in a microwave 3000 digestion system (Anton Paar) for element extraction. Na^+^ and K^+^ concentrations were determined using a flame atomic absorption spectrometer (Analytik Jena).

### Vacuolar pH measurement with BCECF-AM

The pH-sensitive fluorescent dye BCECF-AM was used to measure the vacuolar pH in root cells (Bassil *et al*., 2011b). Four-day-old seedlings grown on vertical plates were collected and incubated in liquid medium containing one-tenth-strength MS medium, 0.5% (w/v) Sucrose, 10 mM MES (pH 5.8), 10 μM BCECF-AM, and 0.02% (v/v) pluronic F-127 (Molecular Probes) for 1 h at 22°CC in darkness. Seedlings were washed once for 10 min before microscopy. Dye fluorescence images were obtained using an LSM880 high-resolution laser confocal microscope (Zeiss, Germany). The fluorophore was excited at 458 and 488 nm, and single emission between 530 and 550 nm was detected for all images. Mature root cells were used for the images. After background correction, integrated pixel intensity was measured at both 458- and 488-nm excitation. Ratio values were used to calculate pH based on a calibration curve using ImageJ software (Bassil *et al*., 2011b). Average ratio values were determined from more than 10 individual seedlings.

### Statistical analysis

All experiments in this study were performed at least three times. Error bars in each graph indicate mean values ± standard error (SE) of replicates. Statistically significant differences between measurements were determined using the independent sample *t*-test (\**P* < 0.05; \*\**P* < 0.01, \*\*\**P* < 0.001) or one-way ANOVA (*P* < 0.05; LSD and Duncan test) in IBM Statistical Product and Service Solutions Statistics software version 24 (IBM, USA).

## Acknowledgments

This work was supported by the National Key R&D Program of China (grant number 2018YFD1000704/2018YFD1000700), the Central Guidance on Local Science and Technology Development Fund of Shandong Province (grant number YDZX2021008), the Agricultural Fine Seed Project of Shandong Province (grant number 2021LZGC006) and the Natural Science Foundation of China (Grant number 31970292, 32170306).

## Author contributions

C.W., C.Z. and X.D. conceived the experiments; S.X. and C.W. write the article; S.X., Y.W., L.Z., Y. B., Y.W., M.L., and J. F. performed experiments; S.T., Y. S., S.Z., J.H., G.Y., and K.Y. provided materials and suggestions. All authors approved the final manuscript.

## Data availability

All relevant data, vectors, and plant materials that support the findings of this study are available from the corresponding author upon request.

## Supporting information

**Figure S1**. Characterization and expression analysis of *SiPLATZ*s in foxtail millet.

**Figure S2**. Transcript levels of *SiPLATZ12* in transgenic foxtail millet.

**Figure S3**. Overexpression of *SiPLATZ12* increases TGW but reduces growth and salt tolerance of transgenic *Arabidopsis*.

**Figure S4**. Relationship between seed size and survival rate with the expression of *PLATZ12* in three types of selected HapHSE and HapHSE^In9^ *Setaria* accessions.

**Figure S5**. Phylogenetic analysis of NHXs (A) and CBLs (B) from foxtail millet (*Setaria italica*) and *Arabidopsis thaliana*.

**Figure S6**. SiPLATZ12 affects expression of salt tolerance related genes in *Arabidopsis*.

**Figure S7**. pH calibration curve and BCECF-AM dye loaded roots.

**Figure S8**. Subcellular localization of SiPLATZ12 in hairy-root of foxtail millet (A) and root tip of transgenic Arabidopsis (B).

**Figure S9**. Diagrams of DNA fragments in the promoter of *SiNHX2, SiCBL4*, and *SiCBL7* in ChIP-qPCR.

**Table S1**. Identification of *PLATZ* genes in foxtail millet.

**Table S2**. Primers and DNA sequences used in this article.

**Table S3**. Identification of *NHX* genes in foxtail millet.

**Dataset EV1**. Accessions used in the haplotype and correlation analysis with TGW, panicle length, main stem diameter, heading date, and SUR.

